# Larger and more dentated hippocampal structure is associated with better memory in the oldest-old

**DOI:** 10.1101/2022.04.10.487750

**Authors:** Vuga Parpura, Hector S Caceres, Sara A Sims, Lawrence Ver Hoef, Anandh K Ramaniharan, Stacy Merritt, Roxanne F Rezaei, Pradyumna K Bharadwaj, Mary Kate Franchetti, David A Raichlen, Courtney J Jessup, Alex G Hishaw, Emily J Van Etten, Theodore P Trouard, David S Geldmacher, Virginia G Wadley, Noam Alperin, Eric S Porges, Adam J Woods, Ron A Cohen, Bonnie E Levin, Tatjana Rundek, Gene E Alexander, Kristina M Visscher

**Affiliations:** University of Alabama at Birmingham Department of Neurobiology; University of Alabama at Birmingham School of Medicine and Evelyn F. McKnight Brain Institute, Birmingham, AL; University of Alabama at Birmingham Neurology Department, Epilepsy Division; University of Miami Miller School of Medicine and Evelyn F. McKnight Brain Institute, Miami, FL; University of Florida and Evelyn F. and William L. McKnight Brain Institute, Gainesville, FL; University of Arizona and Evelyn F. McKnight Brain Institute, Tucson, AZ; University of Southern California, Los Angeles, CA

**Keywords:** hippocampal dentation, cognitive aging, episodic memory

## Abstract

Episodic memory is widely recognized as a critically important aspect of cognition that is often impacted by cognitive and brain aging. Prior work has shown that episodic memory is related to the presence of teeth-like folds on the dentate gyrus, called dentation. We hypothesized that episodic memory performance relates to overall hippocampal structure (i.e., dentation and volume) in an oldest-old cohort. We used data from the McKnight Brain Aging Registry, which consisted of cognitively healthy 85+-year-old adults. We conducted a canonical correlation analysis on 111 participants between a set of episodic memory tests and a set of characterizations of hippocampal structure. The analysis yielded a strong canonical correlation between episodic memory and hippocampal structure (r = 0.491, p = <0.001). The results suggest there is a connection between hippocampal morphology and function in the oldest-old. Our findings suggest that dentation may play an important role in relation to the individual differences observed in episodic memory performance among the oldest old and that hippocampal structure supports healthy cognitive aging.

**Highlights:**

- We characterized hippocampal dentation in a healthy oldest-old sample.
- Hippocampal structure is related to episodic memory in healthy oldest-old adults.
- Memory functioning is related to both hippocampal volume and dentation.

## 1. Introduction

Understanding those factors which lead to healthy cognitive aging is an increasingly essential goal for society as the portion of the population over 85 increases. The 2010 U.S. Census estimated that by 2030, 2.3% of the population will be part of the oldest-old cohort (age 85 and older); by 2050, the number of people 85 years of age and older will further increase to 4.3% of the population (Vincent and Velkoff, 2010). Memory, particularly episodic memory, is sensitive to age and can decline over a lifetime (Dumas, 2015; Luo and Craik, 2008; Spaan, 2015). The decline of memory performance generally begins during middle age and continues into old age (Davis et al., 2003; Nyberg et al., 2012). Some of this decline may be due to the early onset of pathological aging (e.g., dementia) or may be part of a normal aging process. Older age is also associated with higher variance in recall and memory performance, which has been attributed to memory preservation or impairment in individuals (Davis et al., 2003). As the population continues to age, we need to understand memory maintenance since it is an important aspect of healthy cognitive aging. Therefore, our analysis focuses on members of the oldest-old cohort who have healthy and preserved memory.

The case study of Henry Molaison’s (H.M.) medial temporal lobectomy determined that the hippocampus is an essential brain structure important for episodic memory function (Corkin, 1984; Corkin et al., 1997; Scoville and Milner, 1957). Progressing from case studies of patients like H.M., the hippocampus is now studied directly and extensively using neuroimaging, utilizing various measures (e.g., activity, volume, connectivity, dentation). Both healthy and pathological aging might affect highly plastic regions, like the hippocampus, more than less plastic regions in the brain (Fjell et al., 2014). The hippocampus undergoes microstructural and macrostructural changes throughout the lifespan which is suspected to affect the cognitive processes (such as memory) associated with the different subregions of the hippocampus (Langnes et al., 2020).

Although hippocampal volume decreases with age, some data suggest that mean hippocampal volume is not statistically different between young adult and older adult populations (Lupien et al., 2007). Further, these two populations experience large within-group variability in hippocampal volume (Lupien et al., 2007). A meta-analysis found only an inconsistent and weak relationship between hippocampal volume, corrected for intracranial volume, and episodic memory in adults, indicating that hippocampal size may not be sufficient to explain individual differences in episodic memory as previously hypothesized (Van Petten, 2004). This meta-analysis also found better episodic memory performance in children, adolescents, and young adults with smaller hippocampal volume, challenging the idea that larger hippocampal volume contributes to better episodic memory (Van Petten, 2004). Previous studies have shown both positive (Pohlack et al., 2014) and negative (Chantôme et al., 1999) correlations between hippocampal volume and episodic memory scores in healthy adults. However, more recent longitudinal studies have found that the hippocampus atrophies with age, and this shrinking volume is linked to memory decline (Fraser et al., 2015; Kramer et al., 2007; Murphy et al., 2010; Persson et al., 2012; Pudas et al., 2018). Despite this, hippocampal atrophy alone might not account for age-related memory decline (Fjell et al., 2013; Resnick et al., 2000), as similar rates of atrophy have been observed in individuals with stable memory and individuals with declining memory (Pudas et al., 2018). The variability in results in this field suggests that hippocampal volume might not be a stable or the only hippocampal feature related to episodic memory performance.

Unlike hippocampal volume, dentation of the hippocampus has not been as widely studied, despite the prominence of this structure, and its contribution to the term ‘dentate gyrus.’ Despite the fact that these unique structures are prominent and differ considerably from person to person, relatively little research has examined the importance of dentation to cognitive function. To our knowledge, hippocampal dentation has not been observed in other widely used animal models, such as rats and mice. However, indentations have been observed on the margo dentriculatus of higher-order primates (Naidich et al., 2009). An increased number of folds in the medial temporal and occipital lobes of the brain has been positively correlated to intelligence, supporting the argument that more folding, and therefore increased surface area, might improve cognitive processing (Luders et al., 2008). The hippocampus has “teeth-like” folds on its dentate gyrus called dentation; each individual fold is referred to as a dente, as seen in Fig. 1 (Fleming Beattie et al., 2017). Dentes run from the anterior end to the posterior end of the hippocampus. The degree of hippocampal dentation was first measured using a subjective rating scale and correlated with performance on episodic memory tests in a healthy adult sample (age range 20-57 years old) (Fleming Beattie et al., 2017). Additionally, the relationship between hippocampal dentation and hippocampal volume was not significant (Fleming Beattie et al., 2017). A more recent study examined dentation and episodic memory performance in adults with healthy hippocampi and in adults with hippocampi lesioned due to temporal lobe epilepsy (age = 36.64 11.28) In lesioned hippocampi, dentation positively correlated with hippocampal volume; dentation and volume trended toward significance in healthy hippocampi (Kilpattu Ramaniharan et al., 2022). Until now, hippocampal dentation and episodic memory have not been studied in the oldest-old adult cohort.

**Figure 1.**
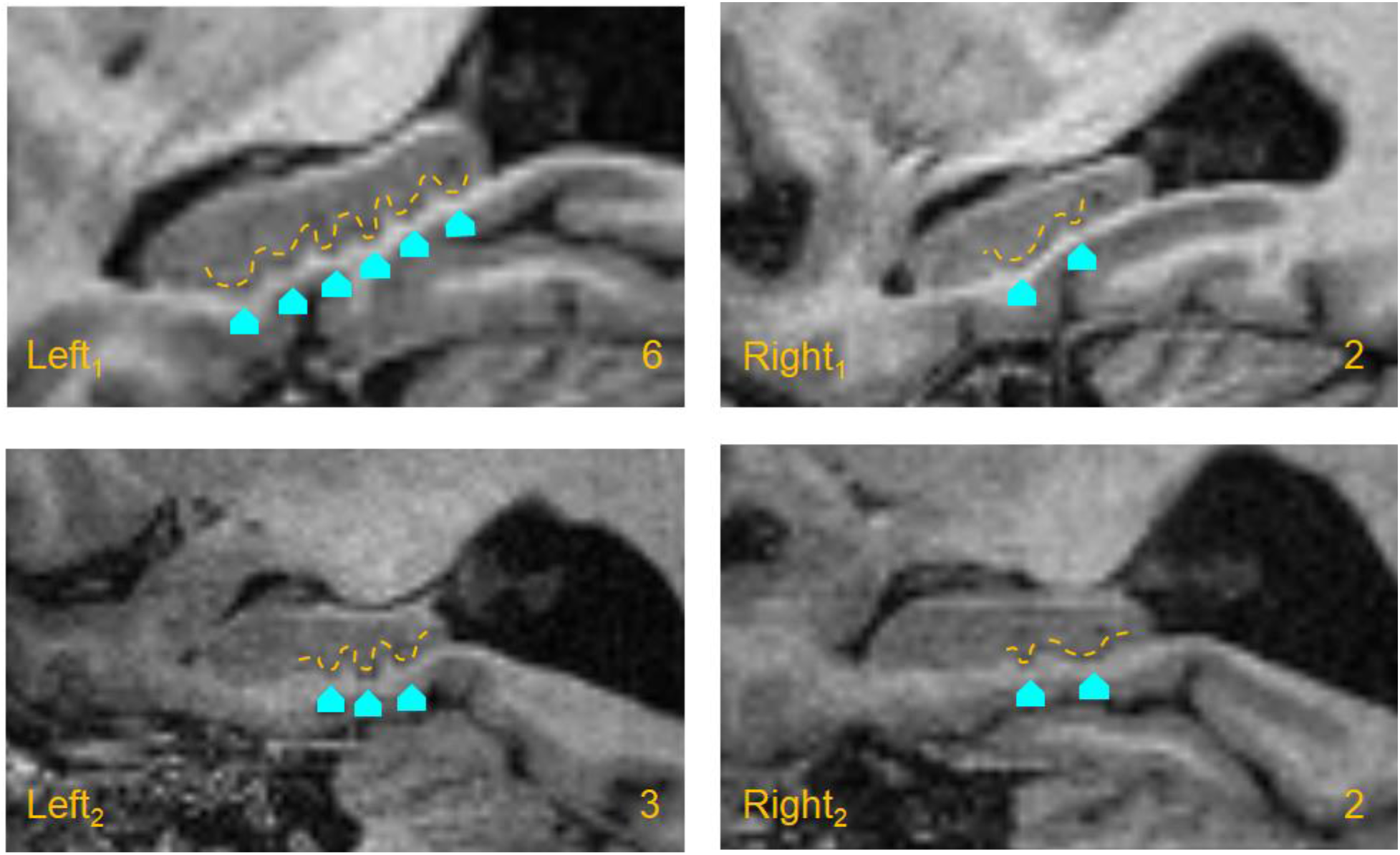
Dentation of the hippocampus. The figure shows anatomical images with a sagittal view of 4 hippocampi. The aquamarine blue arrows point to each individual dente. A dashed yellow line outlines the general shape of each hippocampus and the respective dentes of each hippocampus. Two participants (both age 85) from our sample are shown above. The two top images show the left (Left 1) and right (Right 1) hippocampal dentation counts for one participant in yellow, which were 6 and 2, respectively. The two bottom images show the left (Left 2) and right (Right 2) hippocampal dentation count for a second participant in yellow, which were 3 and 2, respectively.

In this study, we aimed to better understand the link between the morphology of the hippocampus and memory functioning. By studying the hippocampus in the context of a healthy oldest-old sample, we both expand our understanding of memory in this increasingly important age group, and we also take advantage of this population’s large variability in brain structure and in memory performance. We examine hippocampal structure in a novel way by incorporating hippocampal dentation and volume into one hippocampal structure metric. We hypothesized that episodic memory performance is related to multiple features of hippocampal structure in a healthy oldest-old cohort. Therefore, we predicted that better episodic memory performance is related to a larger and more dentated hippocampal structure in a healthy oldest-old cohort.

## 2. Methods

### 2.1 Participants

McKnight Brain Aging Registry (MBAR) participant data was analyzed in this study. MBAR consists of a sample of healthy oldest old (85 years of age and older)individuals. Participants were recruited across four sites: The University of Alabama at Birmingham, The University of Arizona, The University of Florida, and The University of Miami. Participants were recruited via physician referrals, flyers, and mailings. All participants underwent stringent inclusion and exclusion criteria that defined successful physical and cognitive aging: participants were 85 years of age or older, had above a sixth-grade reading level, had no major physical disabilities, were independent in basic and instrumental activities in day-to-day life,were determined to be cognitively unimpaired, and did not have active substance abuse. The participants were screened for neurological disorders, psychiatric disorders, and mild cognitive impairment (MCI), using the Telephone Interview for Cognitive Status (TICS) and the Montreal Cognitive Assessment (MoCA) (Brandt et al., 1988; Nasreddine et al., 2005). Participants also underwent a neurological exam conducted by a neurologist. Figure 2 illustrates the cognitive screening measures used to determine healthy cognition in our oldest-old sample. Data were quality checked for the completion of all four behavioral tests used in our analysis and MRI of sufficient quality to identify the hippocampus. 111 from the total MBAR participants met these criteria and were analyzed for this paper. The average age of our sample was 88.22 years (SD = 3.31 years, Range = 85-99); further description of the sample can be found in Table 1.

**Figure 2.**
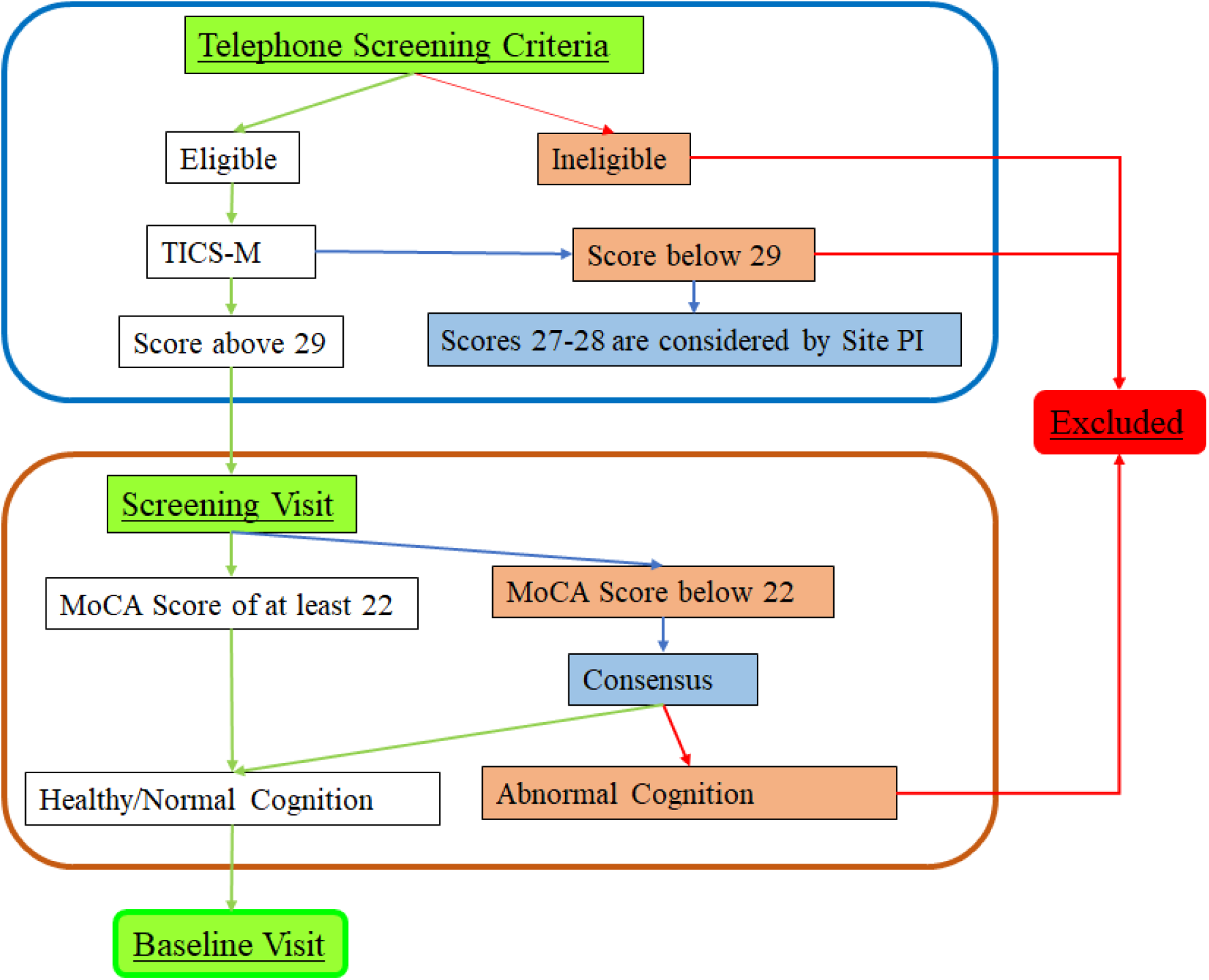
Participant Exclusion/Inclusion Flowchart. The figure above illustrates the order of administration of cognitive screening measures and the process used to determine healthy and normal cognition for our sample of oldest-old adults.

**Table 1.**
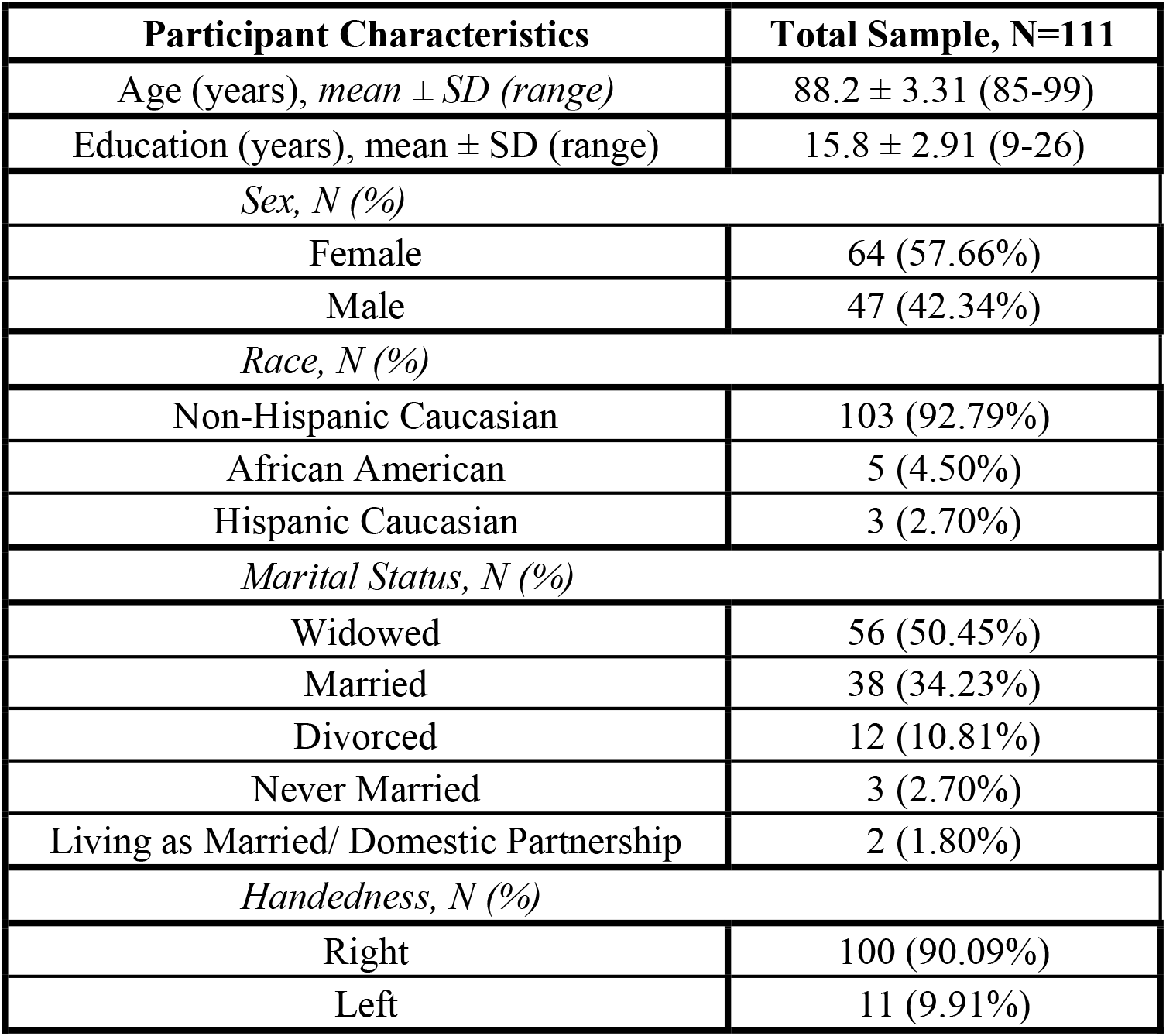
Participants Table. The table above shows the demographic characteristics of our sample of 111 participants included in our analysis.

### 2.2 Episodic Memory Measures

The episodic memory tests used in this analysis were part of a larger test battery that was administered over the course of two separate visits. All behavioral tests were administered by trained and certified testers. All data collected underwent double data entry in Redcap for quality control of discrepancies. Further quality control was done by visually inspecting the data and examining outliers and errors in the data.

#### 2.2.1 MST

The Mnemonic Similarity Test (MST) is an episodic memory test of pattern separation. As part of the MST encoding phase, participants were introduced to a set of images (128 images, 2s each) for which they classified as belonging inside or outside. In the test phase, the participants were shown another set of images (192 images, 2s each). One third of the images shown were repeated images from the previous set (old); one third were images not previously seen during the encoding phase (new); and one third of the images were similar to the previous set, but not exactly the same as the images in the encoding phase (similar) (Stark et al., 2013). The participants classified these images as old, new, or similar to the previous set. The MST, also previously called the BPS-O score, was calculated as the difference between the rate of “Similar” responses given to similar images minus the rate of “Similar” responses given to old images. This is subtracted to account for response bias. A negative score indicates a large number of “similar” responses given for old images. The MST score was used for the analysis.

#### 2.2.2 CVLT

The California Verbal Learning Test Second Edition (CVLT-II) has been widely used as a measure of verbal episodic memory. The CVLT-II was administered to MBAR participants following the guidelines in the CVLT Second Edition Manual (Delis et al., 2000). In the CVLT, participants were given a 16-word list with four semantic categories to learn across five trials. They were then presented with a separate distractor list with four semantic categories different from the first list. The distractor list was immediately recalled. Then, the participants were tested for recall of the original list and for the semantic categories of the list after a short delay (5 minutes). During a long delay (20 minutes), the participants completed different nonverbal tests. Participants were tested for recall of the original list and the semantic categories after the long delay. The Long Delay Free Recall (LFDR) raw scores were used for this analysis.

#### 2.2.3 Benson Figure Complex

The Benson Figure Complex is a visuospatial episodic memory task and is in the neuropsychological battery of the Uniform Data Set from the National Institute on Aging Alzheimer Disease Centers (Morris et al., 2006). Participants were shown the Benson Figure Complex and were instructed to copy the figure. The figure copy was scored for the accuracy and location of elements of the figure. Participants were asked to remember the figure for later recall. During a long delay (10-15 minutes), the participants completed verbal tasks. After the long delay, participants were asked to recall the figure that was copied earlier and draw it from memory, and their drawing was scored using the same criteria as the copy condition. The Benson Delayed score was used in this analysis.

#### 2.2.4 Craft Story 21

The Craft Story 21 is a verbal episodic memory task in the neuropsychological battery of the Uniform Data Set from the National Institute on Aging Alzheimer Disease Centers (Morris et al., 2006). During this task, participants were instructed to listen to a story and remember as much as possible. Immediately after the story, participants were asked to recall the story. Participant responses were scored for both verbatim and paraphrasing recall of story units. During a long delay (20 minutes), the participants completed non-verbal tasks. After the long delay, participants were asked to recall the story and were again scored for both verbatim and paraphrasing recall of story units. If participants did not recall a story, they were prompted with a verbal cue. The Verbatim Delayed Recall score was used in this analysis.

### 2.3 MRI Acquisition

The T1-weighted anatomical scans were acquired using three 3.0 T Siemens Prisma scanners and one 3.0 Skyra Siemens scanner across the four sites with the following parameters: a repetition time (TR) of 2530ms, an echo time (TE) of 3.37m, a field of view [FOV (ap,fh,rl)] of 240 × 256 × 176 mm, a slice gap of 0, a voxel size of 1.0 × 1.0 × 1.0 mm isotropic, and a flip angle (FA) of 7°). All data were harmonized across sites and underwent quality control and visual inspection.

### 2.4 Hippocampal Measures

#### 2.4.1 Hippocampal Dentation Scoring

T-1 weighted MRI images were used to assign a score for hippocampal dentation by viewing the structure of the hippocampus in three dimensions (Fleming Beattie et al., 2017). Although high-resolution multispectral imaging using a combination of T1 and high-resolution T2 scans has been recommended for accurate scoring and to identify more detailed features of dentation, the use of only T1 scans has been suggested for determining counts of dentation with visual ratings along the hippocampus (Fleming Beattie et al., 2017), (Kilpattu Ramaniharan et al., 2022). The subjective dentation score from Kilpattu Ramaniharan et al. (2022) was done for each participant by counting the number of hippocampal dentes in both the left and right hemispheres. The program 3D Slicer was used to view the T-1 weighted images in the sagittal plane (Fedorov et al., 2012). Hippocampal dentation was scored by two trained graders. The rating scale consisted of counting the number of hippocampal dentes of the hippocampus in each hemisphere. The reliability and consistency between the two scorers were determined by generating ICC values (see Results 3.1) for the dentation scores of 111 participants (222 hippocampi in total). The simplified dentation scoring method is shown in Fig. 1. We conducted a paired T-test to determine whether the left and right hippocampal dentation were statistically different.

#### 2.4.2 Hippocampal volume

Hippocampal volumes (mm^3^) were estimated using the Freesurfer Analysis Pipeline (Dale et al., 1999). As done by Fleming Beattie et al. (2017), the anatomical scans were segmented in order to obtain volumes of both left and right hippocampi. The automated hippocampal volume segmentation was visualized and examined for accuracy. We conducted a paired T-test to determine whether the left and right hippocampal volumes were statistically different. Because we are interested in features of the hippocampus in our analysis, we did not normalize hippocampal volumes based on intracranial volume (ICV). Potential atrophy in the whole brain that occurs with age might distort the relationship between volume and dentation when hippocampal volume is corrected for ICV. However, in order to control for the possibility that ICV influences the relationships we observe here, we did perform correlations with intracranial volumes, as described below.

### 2.5 Statistical Analyses

#### 2.5.1 A priori Power Analysis

An *a priori* power analysis was done to determine the sample size needed to detect an effect for our proposed structural equation model. The minimum sample size for the model structure was calculated to be 100, with a power of 0.8, a small effect size (0.1-0.3) (Cohen, 1992), 2 latent variables, 8 observed variables, and an alpha threshold of 0.05.

#### 2.5.2 Canonical Correlation

Canonical correlation analysis (CCA) is a classic multivariate technique used to find linear relationships between two sets of variables (Rosa et al., 2015). The canonical loadings reflect the variance the observed variable shares with the canonical variable. Any canonical loading above |0.30| is considered an important contributing variable to the function (Lambert and Durand, 1975). We used canonical loadings to determine the relative importance of each variable to the function (Kabir et al., 2014; Lambert and Durand, 1975). In order to study the relationship between the episodic memory tests and hippocampal structures, a regularized canonical correlation analysis was conducted (Fig. 3). A partial correlation was performed between the latent variables canonical scores using ICV as a covariate in order to account for total head volume.

**Figure 3.**
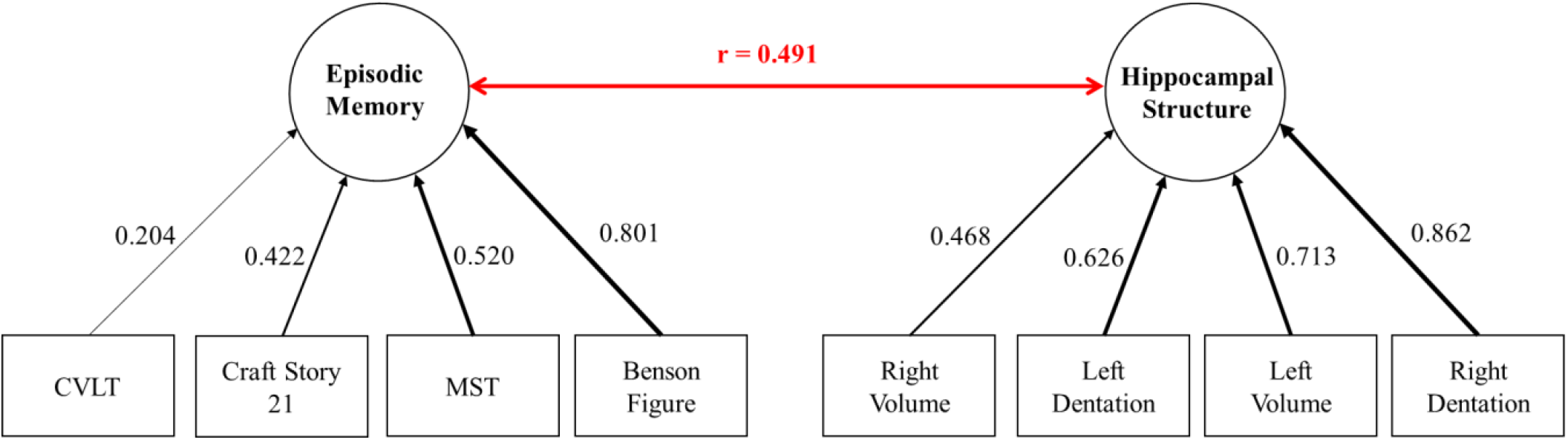
Canonical Correlation Model. The diagram above represents the output of the CCA. The Episodic Memory measures and Hippocampal Structure measures are sorted, left to right, by canonical loadings contributing to two latent variables (“Episodic Memory” and “Hippocampal Structure”). The red arrow represents the canonical correlation (r = 0.491) is illustrated between Episodic Memory and Hippocampal Structure. All of the canonical loadings were originally negative values but were changed to positive values for simplification and interpretability (see section 3.3 for further explanation).

#### 2.5.3 Pearson Correlations

As follow-up descriptive analyses, Pearson correlations were computed between episodic test measures and both hippocampal dentation and volume. This analysis was conducted in order to study the individual effects of hippocampal dentation and volume on episodic test measures.

Previous dentation studies in a healthy adult sample have shown relationships between episodic memory test scores and hippocampal dentation (Fleming Beattie et al., 2017). Correlations between Hippocampal Structure measures and ICV were performed in order to see the effect of ICV on individual measures.

## 3. Results

### 3.1 Hippocampal Dentation ICC

ICC values were calculated to determine the agreement between the dentation assessments of the two trained scorers. The ICC value for the left dentation assessment was r=0.7934 and the ICC value for the right dentation assessment was r = 0.811. Since an ICC value above 0.75 is considered to be excellent (Shrout and Fleiss, 1979), our hippocampal dentation scores are considered reliable and consistent for our analysis.

### 3.2 Observed Variables

The average, standard deviation, and range were calculated for each observed variable of the CCA, shown in Table 2. Two paired samples t-test was done to compare dentation and volume between the left hippocampus and the right hippocampus. The difference in dentation between the left (M= 3.1, SD= 1.3) and right (M= 2.8, SD= 1.5) hippocampus was statistically significant (t(110)= 1.98, p (two-tailed)= 0.0079). However, there was no statistically significant difference (t(110)= 1.98, p (two-tailed)= 0.704)between left (M= 3131.7 mm^3^, SD= 388.2 mm^3^) and right (M= 3142.0 mm^3^, SD= 416.5 mm^3^) volume.

**Table 2.** Episodic Memory and Hippocampal Structure Measures. The table above shows the average scores, standard deviations, and ranges for each measure inputted (i.e., observable variables) into our CCA. Paired Sample T-tests were run for Dentation and Volume. Left Dentation is statistically different from Right Dentation. Left and Right Volumes were not statistically different. The * indicates measures that underwent a paired samples t-test to determine hemispheric statistical significance. The difference in left and right dentation was statistically significant. However, there was no statistically significant difference between left and right volume.

| Observed Variable | Average | SD | Range |
| --- | --- | --- | --- |
| Craft Story 21 | 13.3 | 4.7 | 2 - 24 |
| Benson Figure Complex | 9.7 | 3.3 | 0 - 17 |
| CVLT LDFR | 8.1 | 3.4 | 0 - 16 |
| MST LDI | 0.1 | 0.2 | -0.37 - 0.63 |
| Left Dentation | 3.1* | 1.3 | 1 - 8 |
| Right Dentation | 2.8* | 1.5 | 0 - 6 |
| Left Volume ($\text{mm}^3$) | 3131.7* | 388.2 | 2001.3 - 4213.3 |
| Right Volume ($\text{mm}^3$) | 3142.0* | 416.5 | 1928.6 - 4249.5 |

### 3.3 Canonical Correlation Analysis

Our model (Fig. 3) of a canonical correlation illustrates the relationship between all episodic memory tests and all hippocampal structures. A canonical correlation generates multiple models and Fig. 3 shows the only significant model output by the analysis. The black arrows represent the weighting of each measure to create the latent variable (“episodic memory” and “hippocampal structure.” We found a relatively strong statistically significant canonical correlation between Episodic Memory and Hippocampal Structure variables (r = 0.491 p <0.001), represented by the red arrow in Fig. 3. This model accounted for 33.5% of the variance in the data.

The model assigned weights (Canonical Loadings) to each of the four episodic memory measures that give rise to the “Episodic Memory” factor, all CLs had the same sign, indicating they all tended to covary. When the sign of all the weights is the same (positive or negative), this shows that they all covary in the same direction, like many other types of analyses (e.g, principal components analyses, factor analyses). In this case, every canonical loading value in the model was negative. To make Fig. 3 easier to interpret, we report the absolute value of those canonical loading values; since every value was negative, the interpretation is the same.

Episodic test measure CVLT had a canonical loading (CL) below |0.30| (Lambert and Durand, 1975), indicating it was not an important contributing factor to the function (CL = 0.204). The Benson Figure test showed a strong contribution to the model (CL = 0.801). Right Dentation also showed a strong contribution to the model (CL = 0.862). Yet, Right Volume showed the lowest contribution of the hippocampal structures (CL = 0.468). The partial correlation accounting for ICV resulted in a slightly statistically significant correlation ( r = 0.487, p = <.001).

### 3.4 Correlational Analyses

Although interpretations of the data are based on the CCA above, for completeness we report the individual correlations among episodic memory test measures, among hippocampal structures, and correlations between episodic memory tests and hippocampal structures. Fig. 4 illustrates the relationships between individual episodic memory tests and individual hippocampal structure measures. Verbal episodic memory tests such as CVLT (r = 0.00, −0.17) and Craft Story 21 (r = 0.05, −0.09) had weak correlations to both left and right hippocampal volume. The Benson Figure episodic memory test had the strongest association with the episodic memory tests to all of the hippocampal structure measures (r = 0.27-0.35). Moderate correlations are seen between hippocampal structure measures (r=0.256-0.308). There are strong correlations between left and right hippocampal volume measures (r = 0.679) and left and right hippocampal dentation measures (r = 0.754).

**Figure 4.**
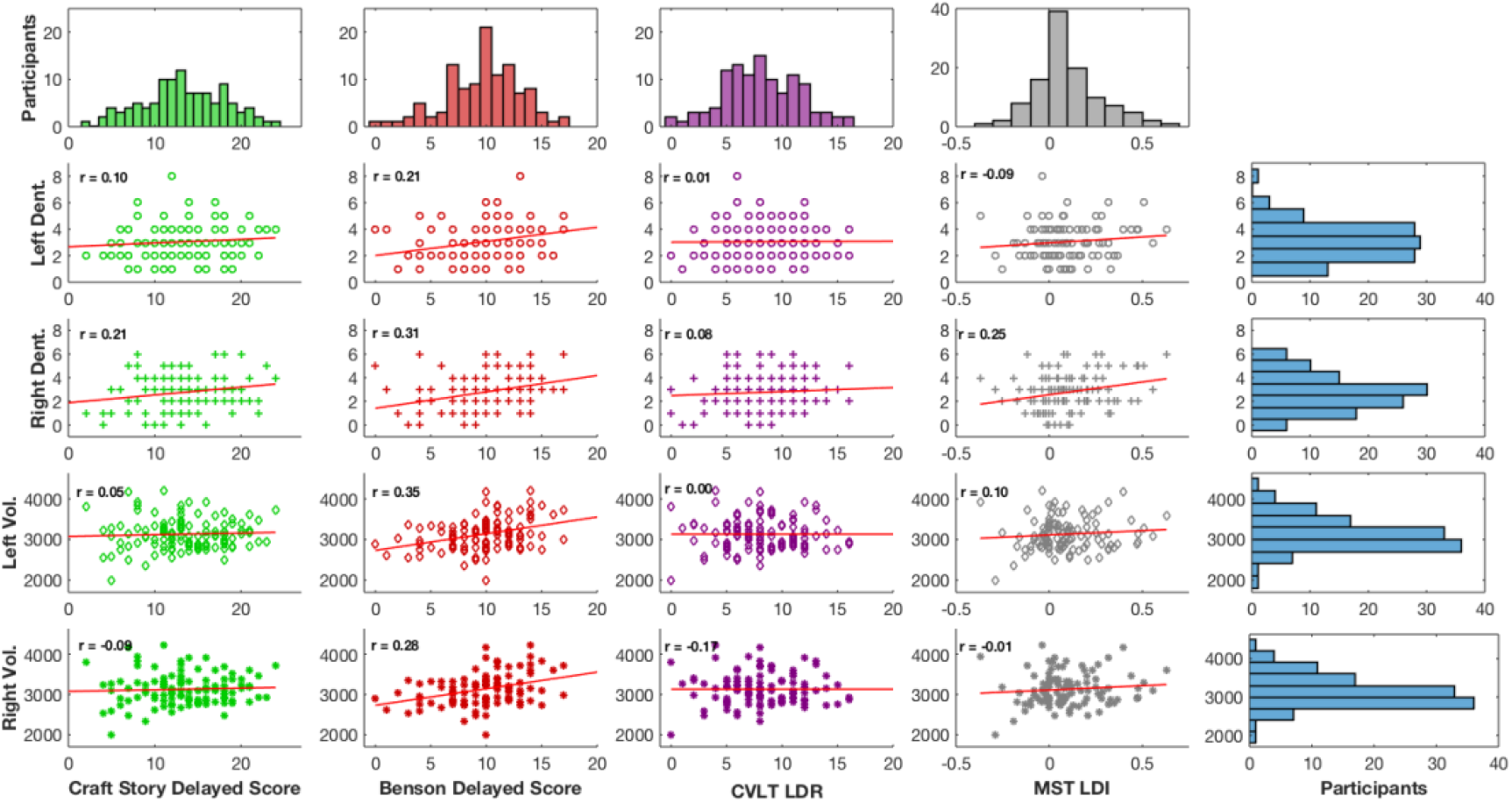
Correlation Plot Diagram. The diagram above represents the correlations between episodic test measures (Benson Figure Complex, Craft Story 21, CVLT, and MST) and hippocampal dentation in each hemisphere (Left Dent= left dentation and Right Dent= right dentation), and hippocampal volume in each hemisphere (Left Vol= left volume and Right Vol= right volume). Colors are representative of episodic memory test measures (green = Craft Story, red = Benson Figure Complex, purple = CVLT, grey = MST). Shapes are representative of hippocampal structures (circles = left dentation, cross = right dentation, diamond = left volume, points = right volume. The r values represent the Pearson correlation value.

**Figure 5.**
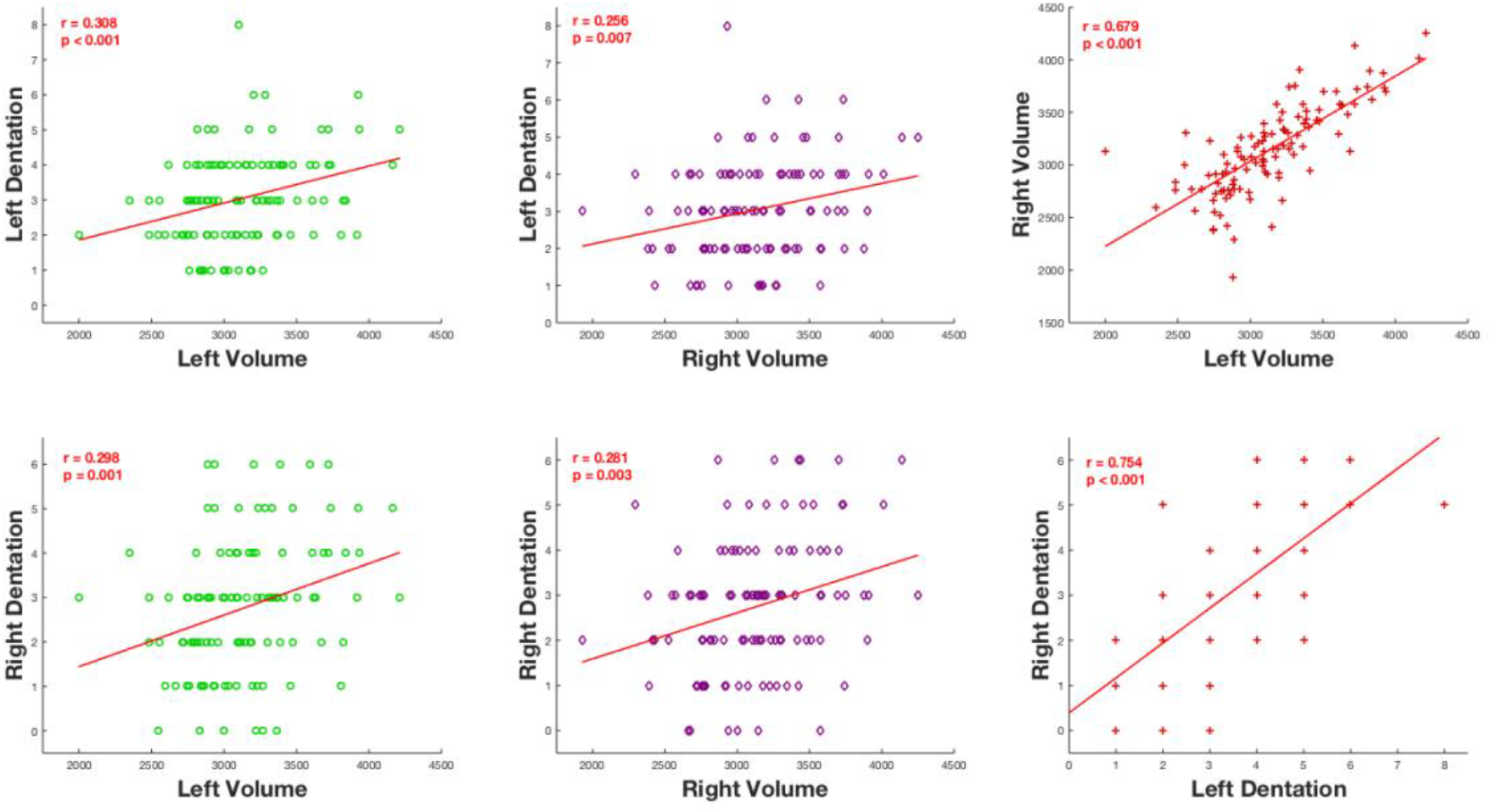
Hippocampal Structures Plot Diagram. The diagram above represents the correlations between the hippocampal structure measures (Left Dentation, Right Dentation, Left Volume, Right Volume). Colors are representative of volume measures (green = Left Volume, purple = Right Volume). Red represents the correlation plots between volume and dentation measures. The r values represent the Pearson correlation values and the p values are representative of the statistical significance.

## 4. Discussion

Our results suggest that both hippocampal dentation and hippocampal volume are important for memory function. A more complex hippocampal structure (ie. higher volume and more dentes) was related to higher episodic memory test measures in this oldest-old adult sample. With the aging of the global population, oldest-old adults will become a greater percentage of the population, making healthy cognitive aging more and more important to understand. Our analysis supports an association between the morphology and memory functioning of the hippocampus in this oldest-old sample, contributing to our understanding of an important aspect of cognitive aging: episodic memory.

### 4.1 Hippocampal Dentation has Variability in Oldest-old Adults

The hippocampus has been central to our understanding of memory. Yet, a connection between specific aspects of morphology, like hippocampal dentation, and function has not been previously demonstrated in this oldest-old adult sample. Fleming Beattie et al. (2017) points to a connection between hippocampal dentation and episodic memory. They found that the degree of hippocampal dentation varied across 22 healthy adults (range: 20-57 years) and was positively associated with episodic memory.

In our findings, we also see this variability in hippocampal dentation in our healthy oldest-old adult group (Table 2). The range of the hippocampal dentation varied between the left and right hemispheres, but the average was similar. Further, the difference in dentation between the left and right hippocampus was statistically significant. Previous literature found no statistical difference in dentation between the left and right hippocampus in a healthy adult sample (Fleming Beattie et al., 2017). The discrepancy in our findings and previous literature may suggest that aging contributes to changes in each hemisphere of the hippocampus. Studies have shown that the hippocampus begins to atrophy in healthy human aging (Fraser et al., 2015). The effects of hippocampal atrophy on hippocampal dentation have not been explored, but our results suggest hippocampal dentation variability occurs in old age. The rates of hippocampal atrophy are similar for those with stable memory and declining memory (Pudas et al., 2018). This finding suggests that hippocampal volume is not the only hippocampal feature related to memory. The variability is seen in our sample, suggesting the previously explored relationship between episodic memory and hippocampal dentation extends into old age. Previous literature points at hippocampal structure not being the only measure related to memory, indicating the variability in dentation seen in the oldest-old sample and hippocampal volume could account for differences in episodic memory. Our findings suggested there was no correlation between hippocampal dentation and intracranial volume (Sup Fig.1). Dentation could then be a useful measure in aging studies as aging causes impacts on brain volume.

### 4.2 The Combination of Hippocampal Volume and Dentation Create a More Complete Measure of Hippocampal Morphology

Previous literature has found conflicting results in the relationship between hippocampal volume and episodic memory (Pudas et al., 2018, Pohlack et al., 2014, Chantôme et al., 1999, Lupien et al., 2007,Van Petten, 2004). The previous results showing a positive relationship between episodic memory and hippocampal dentation seen in healthy adults demonstrated the role dentes may play in memory (Fleming Beattie et al., 2017). Our analysis combined hippocampal volume and hippocampal dentation in order to construct a more comprehensive hippocampal measure. When constructing our latent variable Hippocampal Structure, we divided volume and dentation measures into the left and right hemispheres to account for possible variability between the two hemispheres. As seen in Figure 3, right dentation showed the strongest relationship in the model (CL = 0.862), while right volume showed the weakest relationship of the hippocampal structure measures in the model (CL = 0.468). These findings pointed to right hippocampal dentation as more related to episodic memory. Interestingly, left hippocampal volume showed a stronger relationship in the model (CL = 0.713) than left hippocampal dentation (CL = 0.626), indicating left hippocampal volume contributed more strongly to the Hippocampal Structure. These findings illustrate a potential hemispheric difference in the role the dentation may play on the hippocampal function. Overall, the findings support the idea that all four hippocampal structure measures contribute to our model. The combination of these measures could allow for a more complete measure of hippocampal morphology and point to a stronger connection between morphology and memory functioning in the hippocampus that has not been previously explored.

### 4.3 A Larger and More Complex Hippocampus is Related to Greater Episodic Memory Performance

In the context of healthy aging, better episodic memory performance relates to a larger and more complex hippocampal structure. Fig. 3 illustrates a relatively strong statistically significant canonical correlation (r = 0.491, p < 0.001) between hippocampal structure and episodic memory. CCA determines the variables that have the greatest contribution to the strongest relationship between the two sets. When corrected for ICV using a partial correlation the canonical correlation resulted in similar results (Sup Fig 1., r = 0.487, p < 0.001). The analysis further strengthened the evidence for a positive relationship between hippocampal dentation and episodic memory explored in Fleming Beattie et al. (2017). The latent variable Episodic Memory was composed of different episodic memory tests to include different aspects of episodic memory (pattern separation, verbal, visuospatial) used to create a more complete measure. The latent variable Hippocampal Structure was constructed by both hippocampal volume and dentation in the left and right hemispheres. To our knowledge, our analysis was the first to incorporate both hippocampal dentation and volume as a combined, latent variable of Hippocampal Structure. Our findings point to a connection between the morphology and memory functioning of the hippocampus.

### 4.4 Episodic Memory Measures do not Equally Contribute to Hippocampal Structure

In our analysis, we combined different measures of episodic memory to better measure the domain of episodic memory performance. The model illustrated significant relationships between all of the episodic memory tests except for the CVLT (CL = 0.204). Previous studies have found a positive significant relationship between the CVLT and hippocampal dentation (Fleming Beattie et al., 2017). Yet, previous studies have also found insignificant relationships between hippocampal volume and CVLT (Vuoksimaa et al. 2013). Craft story had the second-lowest contribution to the latent variable (CL = 0.422). Both the CVLT and Craft Story are verbal episodic memory tests. These results show that verbal episodic memory tests had the smallest contribution to the latent variable. Figure 4 showed that both CVLT (r = 0.00; −0.17) and Craft Story (r = 0.05; −0.09) had Pearson correlations near 0 to the hippocampal volume. This along with previous literature is consistent with the notion that hippocampal volume is not related to verbal episodic memory tests (Vuoksimaa et al., 2013). The Benson Figure showed the strongest contribution to the latent variable (CL = 0.801) and strongest Pearson correlations to the hippocampal structures (r = 0.27; 0.31; 0.35; 0.28). The Benson Figure Complex may have more weight in our latent variable since it is specifically aimed at visuospatial episodic memory. The right hippocampus is thought to be involved in the encoding and processing of spatial relationships, thus playing a crucial role in visuospatial memory (Zeidman and Maguire, 2016). The canonical correlation model (Figure 3) showed the strongest relationship to both Benson Figure and Right Dentation. This finding suggests that right dentation could be involved in visuospatial memory. Indicating that hippocampal structure might be related to not only episodic memory functioning but also visuospatial memory.

### 4.5 Limitations and Future Analyses

This work has several limitations. One limitation is the demographically homogenous nature of our participant sample: 92.79% of our sample were Non-Hispanic Caucasians. Further, our sample was also highly educated with an average of 15.8 years of education. This is relevant because research suggests that education influences the impact of cognitive aging on memory (Amieva et al., 2014; Katzman et al., 1989; Rentz et al., 2010; Stern et al., 1995, 1992) such that memory problems occur later in people with more education. Future studies should aim to have more diverse oldest-old samples to more accurately represent the broader population.

Our data were collected across four different sites, each with differences such as test administrators and MR scanners. To mitigate this limitation, prior to data acquisition, personnel at all four sites underwent extensive training, and pilot data acquired at each site were compared. Further, all behavioral data underwent double scoring and double entry; all structural MRI images underwent quality assessment. Our analyses and dataset were not planned or optimized to examine differences among participants of different sex or gender. Future work could control for confounding variables and directly address questions of sex differences. Linear relationships among variables were assumed for statistical analyses. However, we acknowledge the possibility that relationships among variables may not be linear and these assumptions may not be true.

Another limitation of our analysis is the adapted subjective manual scoring of hippocampal dentation. However, we addressed this limitation by running an ICC analysis on the scoring from the two graders. Additionally, the training of the two graders for dentation scoring was given by the neurologist who developed the rating scale in Fleming Beattie et al. (2017). However, Fleming Beattie et al. (2017) notes that merely counting individual dentes alone may not reflect the complexity of the hippocampal structure such as the depth of dentation folds or a curvilinear shape along the length of the hippocampus. A simple dentation count score alone might not most accurately and descriptively represent the structure of hippocampal dentation. The pending development of an objective quantitative method could help mitigate this limitation of subjectivity by more objectively scoring hippocampal dentation (Ramaniharan et al., 2020).

Further study of hippocampal structure across a lifespan could lead to a better understanding of healthy aging, hippocampal structure, and memory. Previous literature found that across the lifespan the hippocampus undergoes micro-and macro-structural changes, which are thought to affect hippocampal function (Langnes et al., 2020). However, Langnes et al. (2020) did not specifically investigate hippocampal dentation. Future work is needed to evaluate how the hippocampal structure, specifically dentation, differently changes over the course of healthy aging in older adult samples and younger adult samples or longitudinally.

## 5. Conclusion

As adult age increases, memory declines, on average (Dumas, 2015; Luo and Craik, 2008; Spaan, 2015). Since memory plays such a vital role in our everyday lives, understanding both memory and its decline in old age is of importance to society. The hippocampus is well known to be important to memory, but details of how its structure supports memory are still being explored. In particular, prominent hippocampal structures, called dentes, give the dentate gyrus its name, but the importance of this structure to the function of the hippocampus is not well understood. This work uses a novel technique to measure hippocampal structure through the use of both dentation and volume. Our study also provides further insight into the range of hippocampal dentation in people with healthy cognition, specifically in an oldest-old study group. Our work supports a relationship between hippocampal morphology and memory functioning in the context of advanced aging. These findings support the idea that hippocampal sub-structure contributes to the memory functioning of the hippocampus. In the oldest-old sample, episodic memory is important to healthy cognitive aging, which our study suggests is predicted by a larger and more complex hippocampus.

## Acknowledgments

We would like to thank everyone who helped with data collection and data management from the MBAR collaborative team. We also thank all of the participants for their contributions to this study. Thank you to other members of the Visscher lab.

## Data Statement

Data is available through the McKnight Brain Aging Registry. The code used for this analysis is available at https://gitlab.rc.uab.edu/hcaceres/hippocampal-dentations/-/blob/master/CorrFigure.m

## Bibliography

Amieva, H., Mokri, H., Le Goff, M., Meillon, C., Jacqmin-Gadda, H., Foubert-Samier, A., Orgogozo, J.-M., Stern, Y., Dartigues, J.-F., 2014. Compensatory mechanisms in higher-educated subjects with Alzheimer’s disease: a study of 20 years of cognitive decline. Brain 137, 1167–1175. doi:10.1093/brain/awu035

Brandt, J., Spencer, M., Folstein, M., 1988. The Telephone Interview for Cognitive Status. Neuropsychiatry Neuropsychol Behav Neurol 1, 111–117.

Chantôme, M., Perruchet, P., Hasboun, D., Dormont, D., Sahel, M., Sourour, N., Zouaoui, A., Marsault, C., Duyme, M., 1999. Is there a negative correlation between explicit memory and hippocampal volume? Neuroimage 10, 589–595. doi:10.1006/nimg.1999.0486

Cohen, J., 1992. A power primer. Psychol. Bull. 112, 155–159. doi:10.1037//0033-2909.112.1.155

Corkin, S., Amaral, D.G., González, R.G., Johnson, K.A., Hyman, B.T., 1997. H. M.’s medial temporal lobe lesion: findings from magnetic resonance imaging. J. Neurosci. 17, 3964–3979.

Corkin, S., 1984. Lasting consequences of bilateral medial temporal lobectomy: clinical course and experimental findings in H.M. Semin. Neurol. 4, 249–259. doi:10.1055/s-2008-1041556

Dale, A.M., Fischl, B., Sereno, M.I., 1999. Cortical surface-based analysis. I. Segmentation and surface reconstruction. Neuroimage 9, 179–194. doi:10.1006/nimg.1998.0395

Davis, H.P., Small, S.A., Stern, Y., Mayeux, R., Feldstein, S.N., Keller, F.R., 2003. Acquisition, recall, and forgetting of verbal information in long-term memory by young, middle-aged, and elderly individuals. Cortex 39, 1063–1091. doi:10.1016/s0010-9452(08)70878-5

Delis, D.C., Kramer, J.H., Kaplan, E., Ober, B.A., 2000. California verbal learning test–second edition. Adult version. Manual. The Psychological Corporation: San Antonio, TX.

Dumas, J.A., 2015. What is Normal Cognitive Aging? Evidence from Task-Based Functional Neuroimaging. Curr. Behav. Neurosci. Rep. 2, 256–261. doi:10.1007/s40473-015-0058-x

Fedorov, A., Beichel, R., Kalpathy-Cramer, J., Finet, J., Fillion-Robin, J.-C., Pujol, S., Bauer, C., Jennings, D., Fennessy, F., Sonka, M., Buatti, J., Aylward, S., Miller, J.V., Pieper, S., Kikinis, R., 2012. 3D Slicer as an image computing platform for the Quantitative Imaging Network. Magn. Reson. Imaging 30, 1323–1341. doi:10.1016/j.mri.2012.05.001

Fjell, A.M., McEvoy, L., Holland, D., Dale, A.M., Walhovd, K.B., Alzheimer’s Disease Neuroimaging Initiative, 2013. Brain changes in older adults at very low risk for Alzheimer’s disease. J. Neurosci. 33, 8237–8242. doi:10.1523/JNEUROSCI.5506-12.2013

Fjell, A.M., McEvoy, L., Holland, D., Dale, A.M., Walhovd, K.B., Alzheimer’s Disease Neuroimaging Initiative, 2014. What is normal in normal aging? Effects of aging, amyloid and Alzheimer’s disease on the cerebral cortex and the hippocampus. Prog. Neurobiol. 117, 20–40. doi:10.1016/j.pneurobio.2014.02.004

Fleming Beattie, J., Martin, R.C., Kana, R.K., Deshpande, H., Lee, S., Curé, J., Ver Hoef, L., 2017. Hippocampal dentation: Structural variation and its association with episodic memory in healthy adults. Neuropsychologia 101, 65–75. doi:10.1016/j.neuropsychologia.2017.04.036

Fraser, M.A., Shaw, M.E., Cherbuin, N., 2015. A systematic review and meta-analysis of longitudinal hippocampal atrophy in healthy human ageing. Neuroimage 112, 364–374. doi:10.1016/j.neuroimage.2015.03.035

Kabir, A., Merrill, R.D., Shamim, A.A., Klemn, R.D.W., Labrique, A.B., Christian, P., West, K.P., Nasser, M., 2014. Canonical correlation analysis of infant’s size at birth and maternal factors: a study in rural northwest Bangladesh. PLoS ONE 9, e94243. doi:10.1371/journal.pone.0094243

Katzman, R., Aronson, M., Fuld, P., Kawas, C., Brown, T., Morgenstern, H., Frishman, W., Gidez, L., Eder, H., Ooi, W.L., 1989. Development of dementing illnesses in an 80-year-old volunteer cohort. Ann. Neurol. 25, 317–324. doi:10.1002/ana.410250402

Kilpattu Ramaniharan, A., Zhang, M.W., Selladurai, G., Martin, R., Ver Hoef, L., 2022. Loss of hippocampal dentation in hippocampal sclerosis and its relationship to memory dysfunction. Epilepsia. doi:10.1111/epi.17211

Kramer, J.H., Mungas, D., Reed, B.R., Wetzel, M.E., Burnett, M.M., Miller, B.L., Weiner, M.W., Chui, H.C., 2007. Longitudinal MRI and cognitive change in healthy elderly. Neuropsychology 21, 412–418. doi:10.1037/0894-4105.21.4.412

Lambert, Z.V., Durand, R.M., 1975. Some precautions in using canonical analysis. Journal of Marketing Research 12, 468–475. doi:10.1177/002224377501200411

Langnes, E., Sneve, M.H., Sederevicius, D., Amlien, I.K., Walhovd, K.B., Fjell, A.M., 2020. Anterior and posterior hippocampus macro-and microstructure across the lifespan in relation to memory-A longitudinal study. Hippocampus 30, 678–692. doi:10.1002/hipo.23189

Luders, E., Narr, K.L., Bilder, R.M., Szeszko, P.R., Gurbani, M.N., Hamilton, L., Toga, A.W., Gaser, C., 2008. Mapping the relationship between cortical convolution and intelligence: effects of gender. Cereb. Cortex 18, 2019–2026. doi:10.1093/cercor/bhm227

Luo, L., Craik, F.I.M., 2008. Aging and memory: a cognitive approach. Can. J. Psychiatry 53, 346–353. doi:10.1177/070674370805300603

Lupien, S.J., Evans, A., Lord, C., Miles, J., Pruessner, M., Pike, B., Pruessner, J.C., 2007. Hippocampal volume is as variable in young as in older adults: implications for the notion of hippocampal atrophy in humans. Neuroimage 34, 479–485. doi:10.1016/j.neuroimage.2006.09.041

Morris, J.C., Weintraub, S., Chui, H.C., Cummings, J., Decarli, C., Ferris, S., Foster, N.L., Galasko, D., Graff-Radford, N., Peskind, E.R., Beekly, D., Ramos, E.M., Kukull, W.A., 2006. The Uniform Data Set (UDS): clinical and cognitive variables and descriptive data from Alzheimer Disease Centers. Alzheimer Dis. Assoc. Disord. 20, 210–216. doi:10.1097/01.wad.0000213865.09806.92

Murphy, E.A., Holland, D., Donohue, M., McEvoy, L.K., Hagler, D.J., Dale, A.M., Brewer, J.B., Alzheimer’s Disease Neuroimaging Initiative, 2010. Six-month atrophy in MTL structures is associated with subsequent memory decline in elderly controls. Neuroimage 53, 1310–1317. doi:10.1016/j.neuroimage.2010.07.016

Naidich, T.P., Duvernoy, H.M., Delman, B.N., Sorensen, A.G., Kollias, S.S., Haacke, E.M., 2009. Duvernoy's Atlas of the Human Brain Stem and Cerebellum: High-Field MRI, Surface Anatomy, Internal Structure, Vascularization and 3 D Sectional Anatomy, illustrated ed. Springer Science & Business Media.

Nasreddine, Z.S., Phillips, N.A., Bédirian, V., Charbonneau, S., Whitehead, V., Collin, I., Cummings, J.L., Chertkow, H., 2005. The Montreal Cognitive Assessment, MoCA: a brief screening tool for mild cognitive impairment. J. Am. Geriatr. Soc. 53, 695–699. doi:10.1111/j.1532-5415.2005.53221.x

Nyberg, L., Lövdén, M., Riklund, K., Lindenberger, U., Bäckman, L., 2012. Memory aging and brain maintenance. Trends Cogn Sci (Regul Ed) 16, 292–305. doi:10.1016/j.tics.2012.04.005

Persson, J., Pudas, S., Lind, J., Kauppi, K., Nilsson, L.-G., Nyberg, L., 2012. Longitudinal structure-function correlates in elderly reveal MTL dysfunction with cognitive decline. Cereb. Cortex 22, 2297–2304. doi:10.1093/cercor/bhr306

Pohlack, S.T., Meyer, P., Cacciaglia, R., Liebscher, C., Ridder, S., Flor, H., 2014. Bigger is better! Hippocampal volume and declarative memory performance in healthy young men. Brain Struct. Funct. 219, 255–267. doi:10.1007/s00429-012-0497-z

Pudas, S., Josefsson, M., Rieckmann, A., Nyberg, L., 2018. Longitudinal Evidence for Increased Functional Response in Frontal Cortex for Older Adults with Hippocampal Atrophy and Memory Decline. Cereb. Cortex 28, 936–948. doi:10.1093/cercor/bhw418

Ramaniharan, A.K., Ver Hoef, L., Zhang, M., Martin, R., Parpura, V., Selladurai, G., 2020. An objective method to quantify hippocampal dentation and predict the side of seizure onset in temporal lobe epilepsy.

Rentz, D.M., Locascio, J.J., Becker, J.A., Moran, E.K., Eng, E., Buckner, R.L., Sperling, R.A., Johnson, K.A., 2010. Cognition, reserve, and amyloid deposition in normal aging. Ann. Neurol. 67, 353–364. doi:10.1002/ana.21904

Resnick, S.M., Goldszal, A.F., Davatzikos, C., Golski, S., Kraut, M.A., Metter, E.J., Bryan, R.N., Zonderman, A.B., 2000. One-year age changes in MRI brain volumes in older adults. Cereb. Cortex 10, 464–472. doi:10.1093/cercor/10.5.464

Rosa, M.J., Mehta, M.A., Pich, E.M., Risterucci, C., Zelaya, F., Reinders, A.A.T.S., Williams, S.C.R., Dazzan, P., Doyle, O.M., Marquand, A.F., 2015. Estimating multivariate similarity between neuroimaging datasets with sparse canonical correlation analysis: an application to perfusion imaging. Front. Neurosci. 9, 366. doi:10.3389/fnins.2015.00366

Scoville, W.B., Milner, B., 1957. Loss of recent memory after bilateral hippocampal lesions. J. Neurol. Neurosurg. Psychiatr. 20, 11–21. doi:10.1136/jnnp.20.1.11

Shrout, P.E., Fleiss, J.L., 1979. Intraclass correlations: Uses in assessing rater reliability. Psychol. Bull. 86, 420–428. doi:10.1037/0033-2909.86.2.420

Spaan, P.E.J., 2015. Episodic and semantic memory functioning in very old age: Explanations from executive functioning and processing speed theories. Cogent Psychology 2. doi:10.1080/23311908.2015.1109782

Stark, S.M., Yassa, M.A., Lacy, J.W., Stark, C.E.L., 2013. A task to assess behavioral pattern separation (BPS) in humans: Data from healthy aging and mild cognitive impairment. Neuropsychologia 51, 2442–2449. doi:10.1016/j.neuropsychologia.2012.12.014

Stern, Y., Alexander, G.E., Prohovnik, I., Mayeux, R., 1992. Inverse relationship between education and parietotemporal perfusion deficit in Alzheimer’s disease. Ann. Neurol. 32, 371–375. doi:10.1002/ana.410320311

Stern, Y., Alexander, G.E., Prohovnik, I., Stricks, L., Link, B., Lennon, M.C., Mayeux, R., 1995. Relationship between lifetime occupation and parietal flow: implications for a reserve against Alzheimer’s disease pathology. Neurology 45, 55–60. doi:10.1212/wnl.45.1.55

Van Petten, C., 2004. Relationship between hippocampal volume and memory ability in healthy individuals across the lifespan: review and meta-analysis. Neuropsychologia 42, 1394–1413. doi: 10.1016/j.neuropsychologia.2004.04.006

Vincent, G.K., Velkoff, V.A., 2010. THE NEXT FOUR DECADES The Older Population in the United States: 2010 to 2050. U.S. Census Bureau.

Vuoksimaa, E., Panizzon, M.S., Chen, C.-H., Eyler, L.T., Fennema-Notestine, C., Fiecas, M.J.A., Fischl, B., Franz, C.E., Grant, M.D., Jak, A.J., Lyons, M.J., Neale, M.C., Thompson, W.K., Tsuang, M.T., Xian, H., Dale, A.M., Kremen, W.S., 2013. Cognitive reserve moderates the association between hippocampal volume and episodic memory in middle age. Neuropsychologia 51, 1124–1131. doi:10.1016/j.neuropsychologia.2013.02.022

Zeidman, P., Maguire, E.A., 2016. Anterior hippocampus: the anatomy of perception, imagination and episodic memory. Nat. Rev. Neurosci. 17, 173–182. doi:10.1038/nrn.2015.24

