## Supplemental Figure 1 for "Larger and more dentated hippocampal structure is associated with better memory in the oldest-old": Supplemental Figures.docx

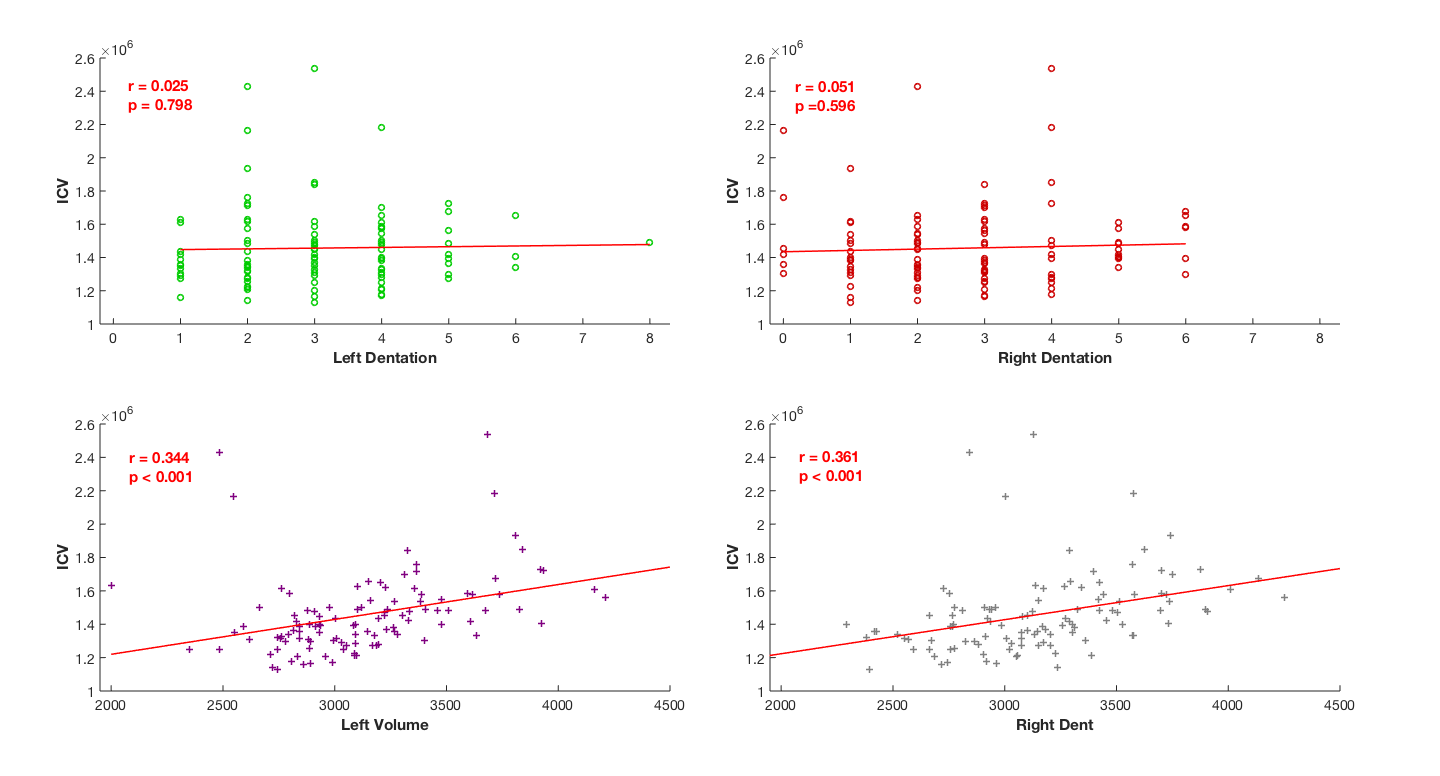

**Supplemental Figure 1. ICV Plot Diagram.** Correlations between intracranial volume (ICV) and Hippocampal Structure measures (left dentation, right dentation, left volume, right volume). r values represent the Pearson correlation values.
